# Neuronal E93 regulates metabolic homeostasis

**DOI:** 10.1101/2022.10.06.511196

**Authors:** Cecilia Yip, Steven Wyler, Shin Yamazaki, Adrian Rothenfluh, Syann Lee, Young-Jai You, Joel Elmquist

## Abstract

Metamorphosis is a transition from growth to reproduction, through which an animal adopts adult behavior and metabolism. Yet the mechanisms underlying the switch is unclear. Here we report that neuronal *E93*, a transcription factor essential for metamorphosis, regulates the adult metabolism and circadian rhythm in *Drosophila melanogaster*. When *E93* is specifically knocked down in neurons, the flies become hyperphagic and obese with increased energy stores and disrupted circadian rhythms. A screen of Gal4 lines targeting subsets of neurons and endocrine cells identified neurons producing GABA and myoinhibitory peptide (MIP) as the main sites of *E93* action. Knockdown of the ecdysone receptor specifically in MIP neurons partly phenocopies the MIP neuron-specific knockdown of *E93* suggesting the steroid signal coordinates adult metabolism via *E93*. The circadian disruption caused by neuronal knockdown of *E93* is also observed when *E93* is knockdown in GABA and MIP neurons. Based on these results we suggest that neuronal *E93* is a key switch for metabolic transition representing an intersectional node between metabolism and circadian biology.

## Introduction

The nervous system integrates environmental and internal cues and generates appropriate physiological and behavioral responses for animals to survive unpredictable and dynamic environments. This physiological and behavioral adaptation often requires a switch between different metabolic states, as best exemplified in insects that go through metamorphosis. Entirely focused on growth, larvae feed constantly to fulfill their metabolic demand, such that their metabolism relies on aerobic glycolysis, similar to cancer metabolism^1-3^. After metamorphosis, however, the focus of an adult switches to reproduction where the adults execute diverse behavioral paradigms to seek mates, succeed in mating, and produce progeny. With these multiple tasks to attend, the importance of feeding diminishes in adults. Indeed, many species of moths, such as *Actias luna*, the American moon moth, and most of the *Saturniidae* family to which *Actias luna* belongs, do not have a feeding organ, and survive and reproduce solely on the nutrients they preserved as larvae^4^.

In *Drosophila*, the switch between growth and metamorphosis is mediated by two essential endocrine signals, Juvenile Hormone (JH) and Ecdysone (20E, a steroid hormone derivative of cholesterol)^5^. JH promotes growth, antagonizing 20E action in metamorphosis, and ensures that larvae grow big enough before they become adults. The balance between these two hormones, which is tightly controlled by the animal’s metabolic state and nutritional environment, determines the timing of development and metamorphosis^6-8^. Metamorphosis itself is a prolonged starvation for over 3 days, and certain metabolically-challenged mutants cannot survive it^1^. In adults, JH and 20E regulate reproduction which also requires an on-off switch upon the females’ mating status, suggesting their continuous roles in balancing metabolic states after metamorphosis.

E93 (*Eip93F*: Ecdysone-induced protein 93F), is one of the master regulators of metamorphosis and has been intensively studied due to its crucial role in the MEKRE93 pathway which many pesticides target^9,10^. E93 belongs to the pipsqueak transcription factor family^11,12^ and contains multiple functional domains such as the DNA binding Helix-Turn-Helix (HTH) domain, the CtBP-interacting domain (CtBP-im) and two potential nuclear hormone interaction domains (NR-box)^13,14^. Conserved from *C. elegans* to mammals^15^, it regulates neurite pruning in *C. elegans*^14^, metamorphosis in insects including *Drosophila*^13^, and is associated with body size variations and lipid metabolism in mammals^16,17^. In *Drosophila*, E93 plays an essential role during metamorphosis as a switch to suppress expression of larval genes and promote expression of adult genes^18^. The enhancer sites targeted by E93 continuously change throughout metamorphosis, suggesting that dynamic interactions between E93 and stage-specific factors allow E93 to serve as a switch for the transition of development stages^19^. Consistent with this role as a switch, E93 is also required for termination of neurogenesis in mushroom body neuroblasts, presumably to help the neurons adapt their adult fate^20^. *E93* expression is maintained at low levels until larva L3 puffstage, the beginning of metamorphosis, when expression is induced by ecdysone (20E)^21^. *E93* expression persists in adults mostly in neurons, including subsets of antennal sense organ and olfactory neurons^22^, suggesting its roles beyond metamorphosis. Despite its essential roles in this developmental program tightly linked to metabolic transition, the mechanisms of E93 action are largely unknown. Moreover, the role of E93 action in metabolism controlled by the nervous system has not been studied.

We report that reduced expression of *E93* in neurons results in hyperphagia and various metabolic and physiological abnormalities. Knockdown of E93 increases attraction to food, food intake, body weight, energy stores and alters circadian rhythm. This failure of metabolic homeostasis is largely due to E93’s action in two populations of neurons: GABA-ergic and myoinhibitory peptidergic (MIP) neurons, both of which have been implicated in controlling food intake, metabolism, circadian rhythm, and sleep^23-29^. MIP neuron-specific knockdown of the ecdysone receptor (EcR), the receptor of ecdysone (20E) which acts upstream of *E93* in metamorphosis, partially phenocopies the MIP neuron-specific knockdown of *E93* in the obese phenotype. Additionally, restoring *E93* only in MIP neurons partially reverses the metabolic abnormalities observed in neuron-specific knockdown of *E93*. Together, our study reveals a novel neuron-specific role of E93 in controlling metabolic homeostasis and circadian rhythm via a steroid hormonal signal.

## Materials and Methods

### *Drosophila* strains and growth conditions

The following lines were obtained from the Bloomington *Drosophila* Stock Center: nSyb-Gal4 (#51635), vGlut-Gal4 (#26160), ChAT-Gal4 (#6793), vGat-Gal4 (#58409), Trh-Gal4 (#38389), Tdc2-Gal4 (#9313), OK107-Gal4 (#854), Akh-Gal4 (#25683), Mip-Gal4 (#51983), Ilp2-Gal4 (#37516), Eth-Gal4 (#51982), Fmrfa-Gal4 (#56837), Lk-Gal4 (#51993), NPF-Gal4 (#25681), phm-Gal4 (#80577), ple-Gal4 (#8848), SifAR-Gal4 (#76670), Tk-Gal4(#51973), Eip93F RNAi (#57868), EcR RNAi (#58286) and vGat RNAi (#41958). E93-RNAi, nSyb-GAL4, mCherry-RNAi lines were outcrossed for five generations to a Berlin background (originally from the Heisenberg lab). Flies were raised on standard cornmeal/molasses food and incubated at 25°C under a 12 h light/12 h dark cycle.

### Quantification of triglycerides (TAG)

Triglycerides and protein were quantified as described by Tennessen *et al*^30^ with minor modifications. 5 flies were decapitated, collected into 1.5 mL tubes, and kept on dry ice before adding 50 μl of 0.5% PBS-T. They were then homogenized, heated for 5 min at 70°C, and centrifuged at 14,000 rpm for three minutes at 4°C. Resulting supernatant was transferred to a new 1.5 mL tube and placed on ice. 200 μl of Infinity Triglyceride reagent (ThermoFisher Scientific, Waltham, MA; catalog no. TR22421) were added to a 96 well plate. 4 μl of supernatant from each sample was transferred to each well in duplicates and the plate was incubated at 37°C for 30-60 minutes. Absorbance levels were read at 540 nm.

### Quantification of protein

Protein level was quantified using the Pierce BCA Protein Assay kit (ThermoFisher Scientific, Waltham, MA; product no. 23228) and used to normalize triacylglycerides (TAG) and glycogen levels. 4 μl of each sample (collected by the method described above) was added to each well of a 96 well plate. Volume of BCA reagent A was determined using the formula: (2 × number of samples + 20) ×100. BCA reagent A and B were mixed (A:B = 50:1) and 100 μl of this mixture was added to each well. The plate was incubated at 37°C for 30 minutes and absorbance values were read at 562 nm of wavelength.

### Quantification of glycogen

Glycogen quantification was performed as described by Tennessen *et al*.^30^ with minor modifications. 2-5 flies per group were decapitated and placed into 1.5 ml tubes before adding 100 μl cold PBS. Samples were homogenized and 10 μl of supernatant were transferred to a new 1.5 ml tube for the protein level measurement. The remaining supernatant was heated at 70°C for 10 minutes and then centrifuged at 14,000 rpm for three minutes at 4°C. 60 μl of resulting supernatant was collected.

Amyloglucosidase digestion solution was prepared by mixing 1.5 μl amyloglucosidase enzyme with 5 ml of 100 mM sodium citrate (pH=5.0). 10 μl of collected supernatant of the glycogen sample was diluted in 90 μl of PBS in a separate 1.5 ml tube. 100 μl of amyloglucosidase solution was added and samples were incubated while shaking at 55°C for one hour. For quantification of glucose, 20 μl of remaining supernatant was added to a 96 well plate in duplicates and were diluted in 60 μl PBS. Similarly, 20 μl of incubated glycogen sample was added to the plate in duplicates and 100 μl prepared GO reagent (o-Dianisidine/Glucose Oxidase/Peroxidase, Sigma Aldrich, St. Louis, MO; GAGO20-1KT) was added to all samples. The plate was incubated at 37°C for 30-60 minutes and 10 μl of 12 N sulfuric acid were added before absorbance was measured at 540 nm^30^.

### Photography

Age-matched control and knockdown flies were imaged using an AM Scope camera (AM Scope, MU1003).

### Quantification of body weight

A group of five age-matched flies was placed in a pre-weighed 1.5 mL tube. Body weight was calculated as mg per fly. Data was collected from 5-10 groups / genotype / sex.

### Quantification of wing morphology

The frequency of unexpanded wings was determined by dividing the number of flies with abnormal wings by the total number of flies in the group. Two to five-day old flies were used.

### Proboscis extension response (PER) assay

Two-day old flies raised on standard media were transferred to vials containing 0.7% agar after 24 h of fasting. A single fly was transferred by use of an oral aspirator and trapped in a 200 µl pipette tip with the head exposed. 1 µl of sucrose solution (150 mM) mixed with 3% erioglaucine (Sigma Aldrich, St. Louis, MO; 861146) was placed on the edge of the pipette tip within range of the proboscis. Proboscis extension and duration of feeding were observed for approximately five minutes.

### Blue dye feeding assay

For each experiment, two-day old flies of 9-16 were fasted for 24 hours on 0.7% agar. To measure baseline food intake without fasting, flies were kept on standard media and allowed to feed *ad libitum*. Feeding vials were prepared by placing a curled strip of filter paper or a pad (Figure 2A) in the center of an empty vial. 400 μl of the blue dye food solution (150 mM sucrose + 3% erioglaucine) was pipetted onto the filter paper. Flies were flipped into the vials and allowed to feed for 5 minutes if previously fasted, and 10 minutes if previously fed *ad libitum*. Flies were immediately incapacitated on dry ice and feeding frequency was determined by dividing the number of flies with blue bellies by the total number of flies. The flies were then individually placed in 1.5 mL tubes, homogenized in 50 μl of water, then centrifuged at 14,000 rpm for 5 minutes. To measure absorbance, 10 μl of supernatant was transferred to a 96 well plate and diluted in 40 μl of water. Absorbance was measured at 630 nm.

### Quantification of proboscis extension

Flies were placed on dry ice to immediately capture their proboscises positions. The three positions of closed, half open, fully open were determined by the degree of proboscis extension.

### RNA isolation and MIP qPCR

RNA was isolated using the Aurum™ Total RNA Mini Kit (Bio-Rad, Hercules, CA) following manufacture instructions. 10 heads were used per replicate. 250 ng of head RNA was converted to cDNA using Superscript IV cDNA Synthesis kit (Thermo Fisher Scientific, Waltham, MA). qPCR was performed using TaqMan probes: *Mip*, Dm02367867_s1 and normalized to *alphaTub* Dm02361072_s1 (Thermo Fisher Scientific). Fold change was calculated using the delta-delta CT (*ΔΔCT*) method. The experiment was repeated twice.

### Starvation assay

Vials were prepared with 0.7% agar. Age-matched males and females were collected and separately placed into starvation vials in 5 groups of 12 flies / vial for a total of 60 flies / sex / genotype. The number of deaths in each group was recorded three times a day until all flies were dead. The assay was performed on five separate cohorts of flies totaling 300 flies / sex / genotype.

### *Drosophila* Activity Monitor (DAM) assay and analyses

Flies were raised on standard media with 12 h light and 12 h dark condition. During the end of the light period, 2-day old flies were placed in the DAM system (TriKinetics Inc., Waltham, MA) where a single fly is loaded in each DAM tube with food on one end. Horizontal locomotor activity was measured by infra-red beam breaks in individual flies under conditions of 12 h light and 12 h dark or in constant darkness for 3 days. Because some nSyb>E93 flies became unhealthy after being kept in the tube more than 5 days, we did not continuously record activity under light and dark condition followed by constant darkness. Naïve 2 days old flies were used for each lighting condition. The average number of beam breaks / 6 min from 16-73 flies of 1-5 cohorts of each genotype and sex were used to generate actograms (ClockLab, Actimetrics, Wilmette, IL). Files that died during 3 days of recording were excluded from the group averaged actogram. Double plotted group averaged actograms were generated from 1-3 cohorts for each genotype and sex with 30 min bin and scaled plot (0-120 counts / 30 min).

### Statistics

We examined whether all experimental groups were normally distributed using Sturges’ class number and a symmetric distribution. All groups appeared to be normally distributed. F test was used to determine equality of variances for two independent groups. Comparisons between control and test groups were carried out using an unpaired two sample *t*-test with unequal variance. Two-way ANOVA was used for Figure 1A and Figure 4B.

**Figure 1:**
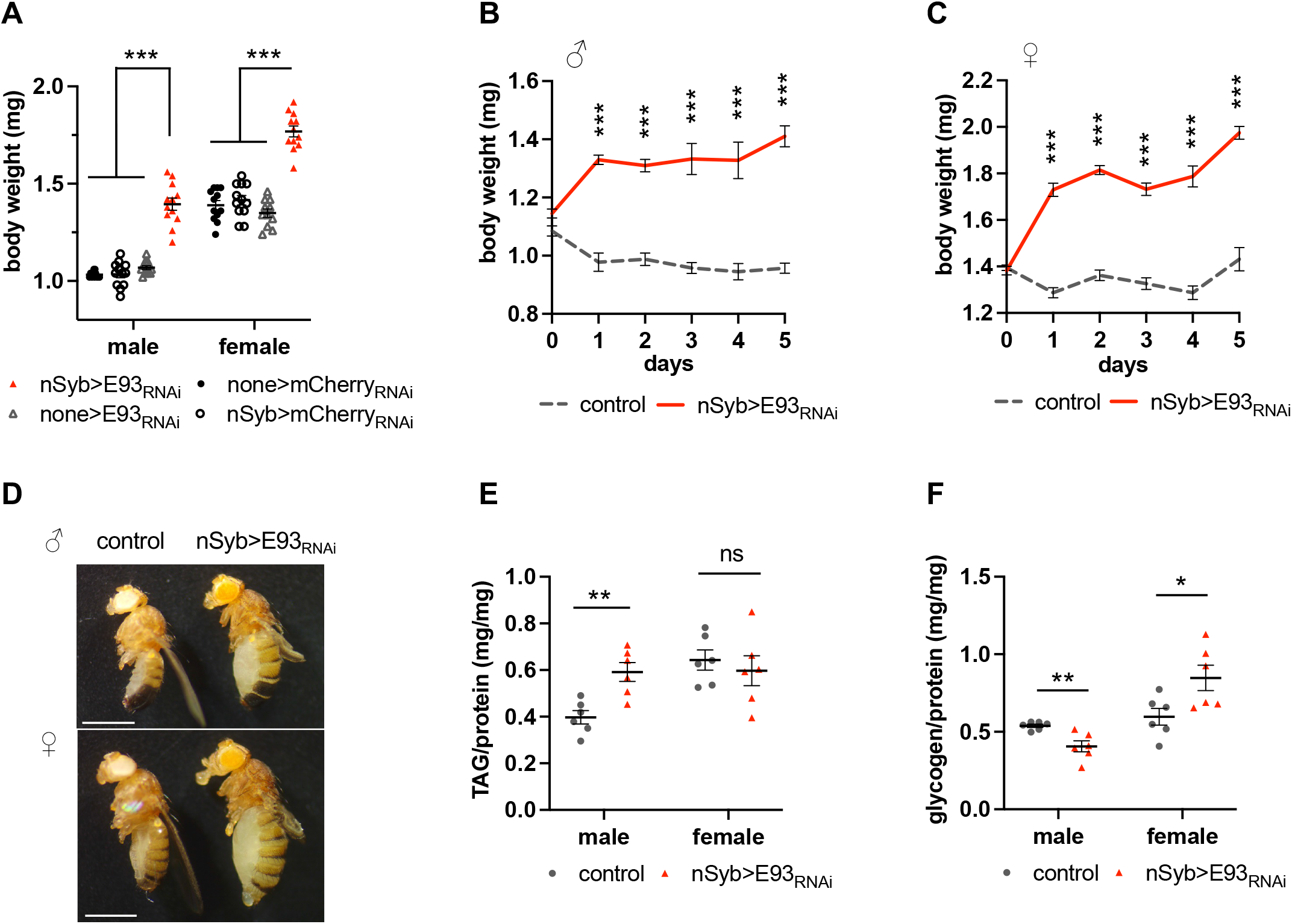
Neuronal specific knockdown of E93 increases body weight and energy stores. **A:** Body weights of (3-day old) of males and females of nSyb>E93_RNAi_ are increased compared to all RNAi only (none>E93_RNAi_ and none>mCherry_RNAi_) and GAL-4 driver (nSyb>mCherry_RNAi_). Each data point represents one group of 5 flies. The bars indicate the mean ± S.E.M. *** *p* < 0.001 by two-way ANOVA. **B-C:** Time-course of body weight measurements in males (B) and females (C) showing a slight (male) or no (female) difference in initial body weights compared to controls (none>E93_RNAi_). Body weights gradually increase in nSyb>E93_RNAi_ flies compared to controls. Errors are S.E.M. *** *p* < 0.001 by Student *t*-test. **D:** Representative images of males (D) and females (E) of control (nSyb>mCherry_RNAi_: white eyes) and nSyb>E93_RNAi_ (orange eyes) flies. Scale bar = 1 mm. **E:** Triglyceride (TAG) levels are increased in nSyb>E93_RNAi_ males but not in nSyb>E93_RNAi_ females when compared to controls (nSyb>mCherry_RNAi_). **F:** Glycogen levels are increased in nSyb>E93_RNAi_ females but not in nSyb>E93_RNAi_ males when compared to controls (nSyb>mCherry_RNAi_). **E-F:** Each data point represents one group of 5 males or one group of 2-5 females. The bars indicate the mean ± S.E.M. * *p* < 0.05, ** *p* < 0.005, ns = not significant by Student *t*-test.

### Ethics statement

No animal work in this study requires specific regulations.

## Data availability

Source data are provided with this paper.

## Results

### Neural specific knockdown of E93 increases body weight and energy stores

When we knocked down the expression of *E93* pan-neuronally using an nSyb-GAL4 driver and a UAS-E93 RNAi (from here, nSyb>E93_RNAi_), the flies became obese; their body weight was increased compared to the three UAS, GAL4, and genetic background control lines (Figure 1A). Because all three controls show very similar body weights, we used either the Berlin K strain carrying UAS-E93 RNAi without any GAL4 driver (none>E93_RNAi_) or nSyb>mCherry_RNAi_ as a control depending on the experiment. nSyb>mCherry_RNAi_, nSyb>E93_RNAi_ and none>E93 _RNAi_ lines were outcrossed to w Berlin for five generations to equalize the background.

The body weight of nSyb>E93_RNAi_ flies gradually increased each day after eclosion (Figures 1B and 1C). Three days after eclosion, both male and female nSyb>E93_RNAi_ flies were visibly obese (Figure 1D). When we measured abdomen width, both knockdown males and females had increased abdomen size (Figure S1A). Compared to controls, female knockdowns had significantly enlarged abdomens beginning one day after eclosion, but both nSyb>E93_RNAi_ and controls retained similar numbers of mature eggs (14.4 ± 5.3, n=18, vs 14.9 ± 4.3, n=13, supplementary methods), indicating that the enlarged abdomen observed in knockdowns was not due to an increased number of retained eggs.

When we measured the levels of triacylglycerides (TAG), the main storage form of lipid, and glycogen, we found that males had more TAG, whereas females had more glycogen than controls (Figures 1E and 1F). These results suggest that neuronal E93 regulates energy stores and that the energy stores may be regulated in a sexually dimorphic manner.

### Neural specific knockdown of E93 increases food intake

Next, we asked whether the increased body weight and energy stores in nSyb>E93_RNAi_ flies were due to an increase in food intake. When nSyb>E93_RNAi_ flies were introduced to a vial with a pad containing sucrose solution (150 mM) and blue dye, a higher percentage of nSyb>E93_RNAi_ flies gathered to the pad than controls (Figure 2A). Consistent with this observation, nSyb>E93_RNAi_ flies showed increased feeding frequency measured by the percentage of total fed flies with blue abdomens (Figure 2B). Furthermore, the majority of nSyb>E93_RNAi_ flies stayed on the pad throughout the experiment, whereas the control flies quickly ate and left within several seconds (Supplementary video 1). The proboscis, the feeding organ, of nSyb>E93_RNAi_ flies was fully extended at a higher frequency in nSyb>E93_RNAi_ flies than in the control, suggesting that E93 knockdown flies are feeding more frequently than the controls (Figures 2C and 2D).

**Figure 2:**
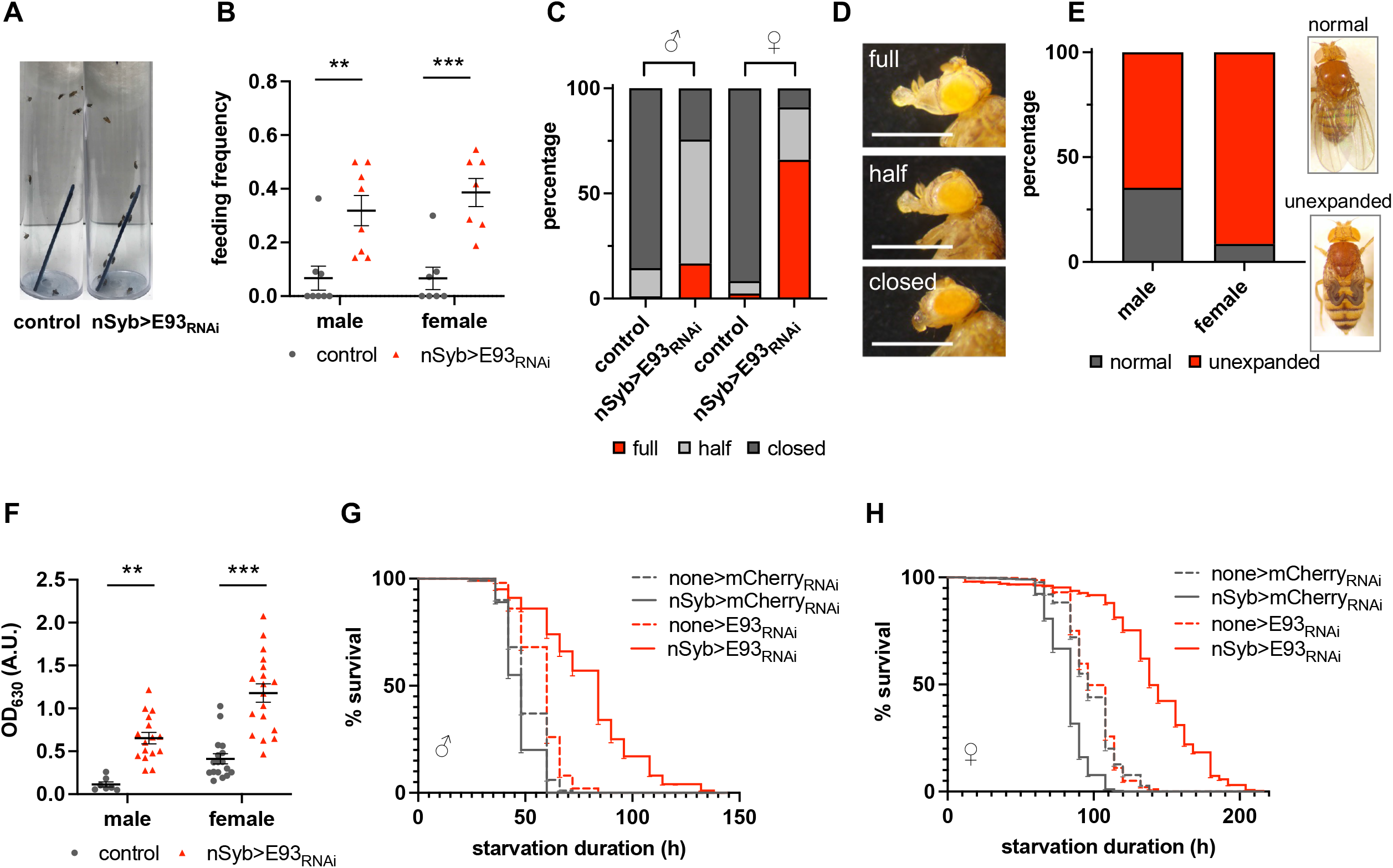
Neuronal specific knockdown of E93 increases feeding frequency, food intake and starvation survival. **A-B:** nSyb>E93_RNAi_ flies are more attracted to food (**A**) and show a higher feeding frequency than the control (nSyb>mCherry_RNAi_) (**B**). Each data point represents one experiment of 9-16 flies. The bars indicate the mean ± S.E.M. **C:** nSyb>E93_RNAi_ flies constantly feed as indirectly measured by % of flies with extended proboscis. N=144-187 males, 83-100 females **D:** Representative images of proboscises in each state of closed, half-open and fully open states. Scale bar = 100 µm **E:** The % of unexpanded wings in nSyb>E93_RNAi_ flies. N= 82 males, 113 females **F:** nSyb>E93_RNAi_ flies have increased food intake. After 10 min of feeding, the flies were homogenized, and the food intake was measured by absorbance of the blue dye at 630 λ. none>E93_RNAi_ flies were used as controls. Each data point represents one sample. The bars indicate the mean ± S.E.M. **G-H:** Both males (G) and females (H) of nSyb>E93_RNAi_ survive starvation better than all three controls. ** *p* < 0.005, *** *p* < 0.001, ns = not significant by Student *t*-test.

The Capillary Feeding (CAFE) assay is commonly used to assess food intake and relies on the ability of flies to reach a suspended food source and hang upside down long enough to feed until satiated^31^. As nSyb>E93_RNAi_ flies have unexpanded wings (Figure 2E), we were unable to perform the CAFE assay reliably. Instead, we modified a blue dye feeding assay and determined the amount of individual food intake by measuring dye absorbance at λ_630_ (see Methods)^32^. Virgin male and female nSyb>E93_RNAi_ flies fasted for 24 h had increased food intake compared to controls (Figure 2F).

We used the proboscis extension reflex (PER) assay to further examine the observed hyperphagia of nSyb>E93_RNAi_ flies in detail. Flies were individually trapped in a pipet tip and presented with 1 µl of sucrose solution containing blue dye under the proboscis^27^. The duration of proboscis extension and the volume of consumed food were measured over 5 min by subtracting the remaining volume from the 1 µl originally presented. Although the number of samples we tested with this method was small, we were able to observe a striking difference between nSyb>E93_RNAi_ flies and controls. All nSyb>E93_RNAi_ flies maintained extended proboscises for the entire 5 min (100%, 5 out of 5) and 80% (4 out of 5) of flies finished 1 µl within 5 min. On the other hand, only 40% (2 out of 5) of the control flies had a single bout of feeding lasting 30 s or less and consumed less than 0.4 µl (Supplementary videos 2, 3). The remaining 60% did not extend their proboscis to feed at all.

In females, changes in midgut size are often associated with increased food intake when nutritional demand is high^33^. The average diameter of the midgut of nSyb>E93_RNAi_ females was increased compared to controls, suggesting an increase in food intake (Figures S1B and 1C). In males increased food intake is often associated with increased defecation rate^34^. nSyb>E93_RNAi_ males show an increased defecation rate compared to controls, suggestive of increased food intake (Figure S1D). Both male and female nSyb>E93_RNAi_ flies survived starvation longer than controls (Figures 2G and 2H), suggesting that their increased energy stores rendered them starvation resistant. Taken together, our data indicate that neuronal E93 is required to control food intake and metabolism.

### E93 acts mainly in GABA-ergic and MIP-producing neurons

To determine the sites of E93 action, we screened 17 GAL4 lines in which the expression of the E93 RNAi construct was limited to subsets of specific classes of neurons and/or endocrine cells. We used glycogen levels in females as a readout, reasoning that the big difference in glycogen levels between nSyb>E93_RNAi_ and the control would provide us a reliable resolution for detection. The 17 lines include the three classical neurotransmitter-producing neurons: vGlut (glutamate), ChAT (acetylcholine) and vGAT (GABA); three monoamine-producing neurons: ple (dopamine), Tdc2 (octopamine/tyramine) and Trh (serotonin); and the mushroom body specific line, OK107, which is important for learning and memory. The remaining ten drivers were selected based on their known roles in feeding and metabolism, and include cells producing MIP (myoinhibitory peptide)^27^, Akh (Adipokinetic hormone)^35^, Eth (Ecdysone triggering hormone)^36^, Phm (ecdysone producing cells of the prothoracic glands)^37^, Tk (Tachykinin)^38,39^, Npf (neuropeptide F)^40^, Fmrfa (FMRFamid neuropeptide), Dilp2 (Insulin-like peptide), Lk (Leucokinin), and SIFaR (SIFamide receptor)^41^. To control for the effect of different GAL4s, we used UAS-mCherry_RNAi_ as a control for each driver. Of note, UAS-mCherry_RNAi_ (none>mCherry_RNAi_) flies had comparable metabolic parameters, including body weight and starvation response, to other controls (Figures 1A, 2G and 2H).

Flies with a knockdown of *E93* in GABA-ergic neurons or in MIP neurons had increased glycogen levels (Figure 3A). Interestingly *E93* knockdown in serotonergic neurons (driven by Trh-GAL4) reduced glycogen levels, suggesting a differential role of E93 depending on neuron types. Glycogen levels remained unaltered in other knockdown lines (Figure 3A).

**Figure 3:**
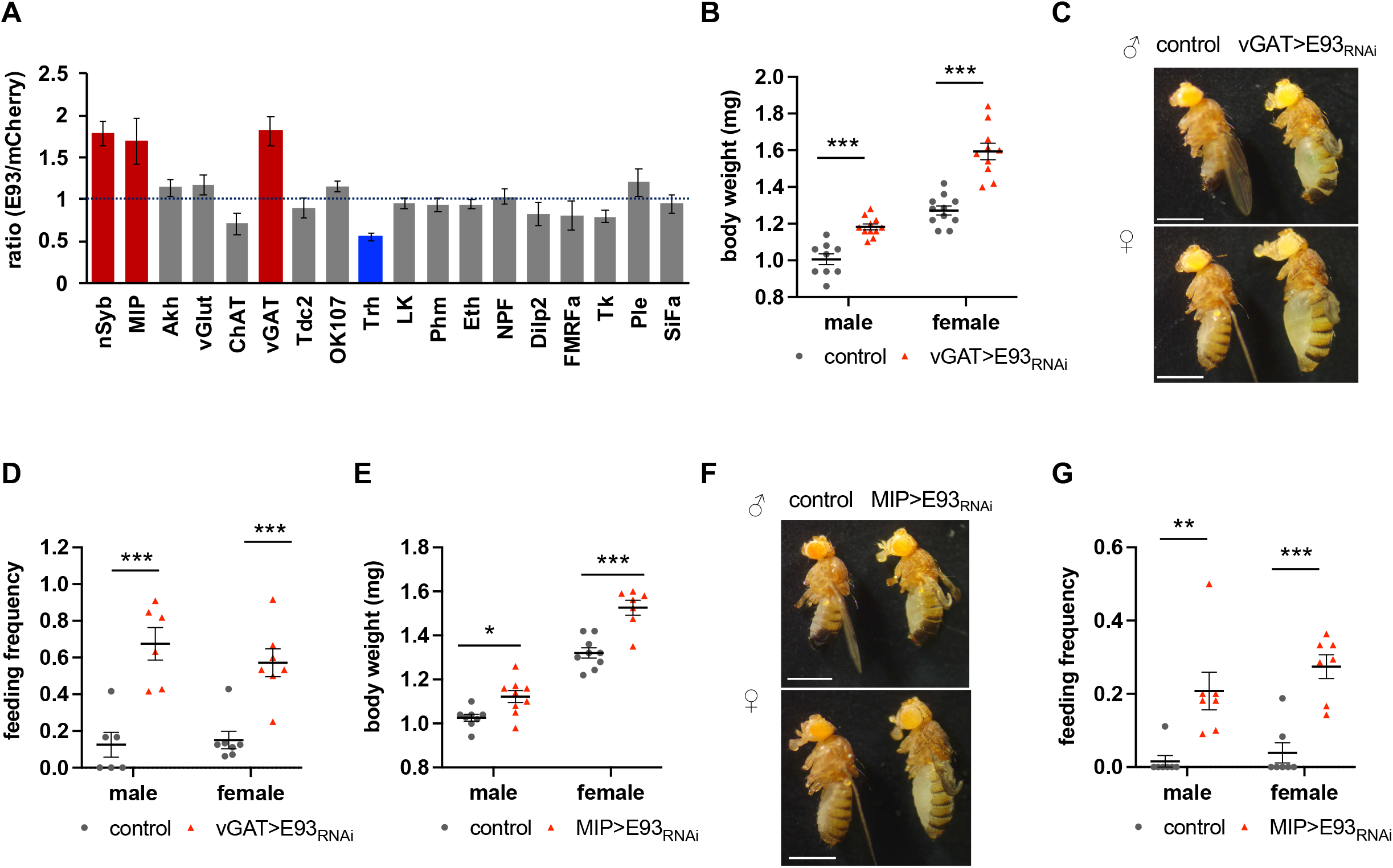
Loss of E93 in GABA and MIP neurons underlie metabolic changes. **A:** Glycogen levels after knockdown of E93 in all neurons and 17 subsets of neurons in females indicates MIP-producing and GABA-ergic neurons most significantly contribute to the increase of glycogen seen in nSyb>E93_RNAi_. **nSyb**: pan-neuronal, **MIP**: Myoinhibitory, **Akh**: Adipokinetic, **vGlut**: Glutamatergic, **ChAT**: Cholinergic, **vGAT**: GABAergic, **Tdc**: Octopaminergic/ tyraminergic, **OK107**: mushroom body, **Trh:** Serotonergic, **LK:** Leucokinin-producing, **Phm:** ecdysone producing cells of the prothoracic glands, **Eth:** Ecdysis triggering hormone-producing, **NPF:** Neuropeptide F-producing, **Dilp2:** Insulin-like peptide-producing, **FMRFa:** FMRFamide peptide-producing, **TK:** Tachykinin-producing, **Ple:** Dopaminergic, **SiFa:** SIFamide-producing. The red-colored bars indicate a significant increase in glycogen level and the blue bar indicates a significant decrease. Data was plotted as the mean ± S.E.M. Each Gal4 driver with mCherry_RNAi_ were used as controls. **B-D:** vGAT>E93_RNAi_ phenocopies nSyb>E93_RNAi_ with increased body weight (B), obese appearance (C) and increased feeding frequency (D). vGAT>mCherry_RNAi_ was used as controls. **B:** Each data point represents one group of 5 flies. **D:** Each data point represents one experiment of 9-16 flies. The bars indicate the mean ± S.E.M. **C:** Scale bar = 1 mm. **E-G:** MIP>E93_RNAi_ phenocopies nSyb>E93_RNAi_ with increased body weight (E), obese appearance (F) and increased feeding frequency (G). MIP>mCherry_RNAi_ flies were used as controls. **E:** Each data point represents one group of 5 flies. **G:** Each data point represents one experiment of 9-16 flies. The bars indicate the mean ± S.E.M. * *p* < 0.05, ** *p* < 0.005, *** *p* < 0.001 by Student *t*-test.

Specific GABA-ergic interneurons are known to play a critical role in inhibiting feeding^23^. Indeed, body weight and feeding frequency were greater in both male and female vGAT>E93_RNAi_ flies compared to controls (Figures 3B-3D). In addition, a higher percentage of vGAT>E93_RNAi_ males and females extended their proboscises than controls (93% of vGAT>E93_RNAi_ (N=57) vs. 2% of vGAT>mCherry_RNAi_ (N=46)). The abdomen size of the vGAT>E93_RNAi_ females was increased without an increase of the number of retained eggs (Figures S2A and S2B). These results validate the use of glycogen levels as a readout to identify the sites of E93 action in regulation of food intake and body weight. Similar to nSyb>E93 flies, vGAT>E93_RNAi_ flies also exhibited a wing phenotype; 50% of males and 82% of females had unexpanded wings while 100% of controls had normal wings (N=32 for males, N=50 for females). These results indicate that GABA-ergic neurons are one of the main sites of E93 action.

Next, we examined flies with *E93* knocked down in MIP neurons. MIP/allatostatin-B homologs are present in almost all animals^42^, and serve versatile roles related to metabolism such as feeding and metamorphosis^43^. In *Drosophila*, MIP is expressed in approximately 70 neurons in the areas important for food sensation such as the antennal lobe and subesophageal zone (SEZ), and regulates mating receptivity, satiety, fat storage, and sleep^27-29^. Both males and females of MIP>E93_RNAi_ are obese and feed more (Figures 3E-3G), indicating MIP neurons are also where E93 acts. The MIP>E93_RNAi_ females recapitulate the previously reported phenotypes of the MIP mutant females^27^. In addition, MIP>E93_RNAi_ females have increased abdomen size without changing in the number of retained mature eggs (Figures S2C and S2D)

Because MIP>E93_RNAi_ flies show similar phenotypes to those of MIP mutants and because E93 is a transcription factor, we next asked whether MIP transcription levels are reduced in nSyb>E93_RNAi_ flies. qPCR from head samples of nSyb>E93_RNAi_ female flies showed reduced levels of MIP mRNA, suggesting that E93 controls food intake and metabolism partly via MIP signaling (Figure 4A).

To confirm that MIP neurons are a site of E93 action, we introduced GAL80, a repressor of GAL4, only in MIP neurons in nSyb>E93_RNAi_ flies. MIP driven GAL80 represses GAL4 driven by nSyb in MIP neurons and thus expression of *E93* is restored only in MIP neurons, while nSyb-GAL4 continues to drive *E93* RNAi in all other neurons. Introducing GAL80 only in MIP neurons partially but significantly reverses the body weight phenotype of nSyb>E93_RNAi_ flies (Figures 4B and 4C), confirming that MIP neurons are one of the sites of E93 action.

**Figure 4:**
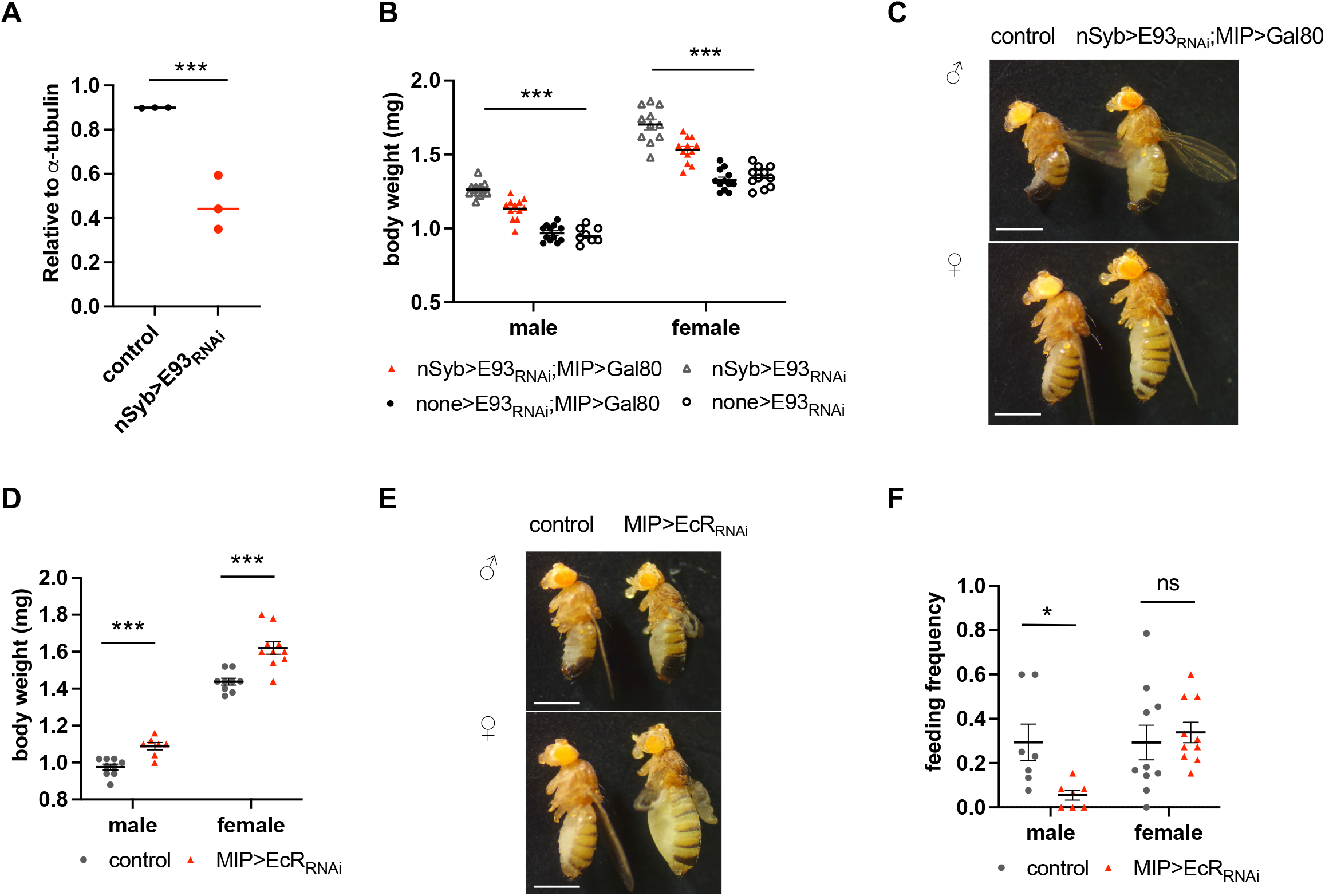
E93 acts in MIP neurons downstream of ecdysone receptor. **A**: MIP mRNA levels are reduced in nSyb>E93_RNAi_ compared to controls (none>E93_RNAi_). Each data point represents 10 female heads. It is a representative graph of two independent experiments. **B-C:** MIP>GAL80 partially but significantly reverses the increased body weight phenotype of nSyb>E93_RNAi_. Each data point represents one group of 5 flies. The bars indicate the mean ± S.E.M. ** *p* < 0.005, *** *p* < 0.001 by Two-way ANOVA. C: Scale bar = 1 mm. **D-F:** MIP>EcR_RNAi_ phenocopies MIP>E93_RNAi_ with increased body weight (D) and obese appearance (E), but not in feeding frequency (F). MIP>mCherry_RNAi_ flies were used as controls. **D:** Each data point represents one group of 5 flies. **F:** Each data point represents one experiment of 9-16 flies. The bars indicate the mean ± S.E.M. * *p* < 0.05, ** *p* < 0.005, *** *p* < 0.001, ns = not significant by Student *t-*test.

### Steroid nuclear receptor controls metabolism in MIP neurons

During the process of metamorphosis, ecdysone binds to the ecdysone receptor (EcR) to induce expression of *E93*^12^. To delineate the signaling pathway of E93, we examined whether EcR also acts upstream of E93 and mediates the metabolic phenotypes observed in adults. Ecdysone and EcR regulate many essential functions not only during metamorphosis but also in adult flies. Ecdysone is secreted mainly from the fat body in males and ovaries in females, and controls reproduction and metabolism^10,44,45^. Pan-neuronal and GABA-ergic neuron-specific knocking down of EcR (nSyb>EcR_RNAi_ or vGAT>EcR_RNAi_) are lethal, thus we could not investigate the role of EcR in feeding and metabolism in these cells. However, using MIP-GAL4 and UAS-EcR_RNAi_ lines, we found that the MIP-specific knockdown of EcR (MIP>EcR_RNAi_) phenocopied MIP>E93_RNAi_. MIP>EcR_RNAi_ flies showed increased body weight and were visibly obese (Figures 4D and 4E), suggesting a genetic interaction of E93 with EcR in MIP neurons and regulates metabolism. Similar to nSyb>EcR_RNAi_ females, MIP>EcR_RNAi_ females had increased abdomen size without changes in the number of retained mature eggs (Figures S3A and S3B). However, we found that the feeding frequency of MIP>EcR_RNAi_ was either not different from the controls (Figure 4F female) or slightly reduced (Figure 4F male). The mechanisms underlying this dissociation between weight gain and food intake are unknown. One possibility is that MIP>EcR_RNAi_ flies have altered patterns of food intake and activity (see below). Alternatively, as EcR plays a significant role from the birth, knockdown of EcR in MIP neurons throughout development could result in adaptations or defects that mask the feeding phenotype seen in adult MIP>E93_RNAi_. Nonetheless, GABA is important for controlling food intake^23,24^ and is secreted from some MIP neurons^46^. We therefore examined whether GABA produced in MIP neurons contributes to the observed nSyb>E93_RNAi_ phenotypes. However, no differences in body weight were observed in flies with knockdown of the vesicular transporter of GABA (vGAT) in MIP neurons, suggesting that GABA released by MIP neurons might not contribute to the obesity phenotype of MIP>E93_RNAi_ flies (Figures S3C and S3D).

### Neuronal knockdown of E93 alters locomotor activity and disrupts circadian rhythm

The reciprocal interaction between biological rhythm and metabolism is conserved in many animals and metabolic abnormality is often associated with abnormal locomotive activity^47,48^. For example, *Clock* mutant mice is obese^49^ and high-fat diet disrupts the activity and feeding rhythms^50,51^. Studies in *Drosophila* also reveal evolutionarily conserved mechanisms of circadian rhythm controlled by metabolism^52-55^. When we measured the daytime and nighttime activity of nSyb>E93_RNAi_ flies using the *Drosophila* Activity Monitoring (DAM) system, both females and males showed overall reduced activity under 12 h light and 12 h dark condition compared to the controls (Figure 5A). Similar reductions in activity were observed in MIP>E93_RNAi_ flies (Figure 5C). However, only a modest reduction in activity was observed in MIP>EcR_RNAi_ flies and no obvious change in activity was observed in vGAT>E93_RNAi_ flies (Figures 5D, 5B). Interestingly, the nighttime activity of vGAT>E93_RNAi_ was elevated (Figure 5B). GABA activity is required to promote sleep by suppressing wake-promoting neurons such as pigment dispersing factor (PDF) producing neurons and monoaminergic neurons such as dopamine or octopamine^26,56^.

**Figure 5:**
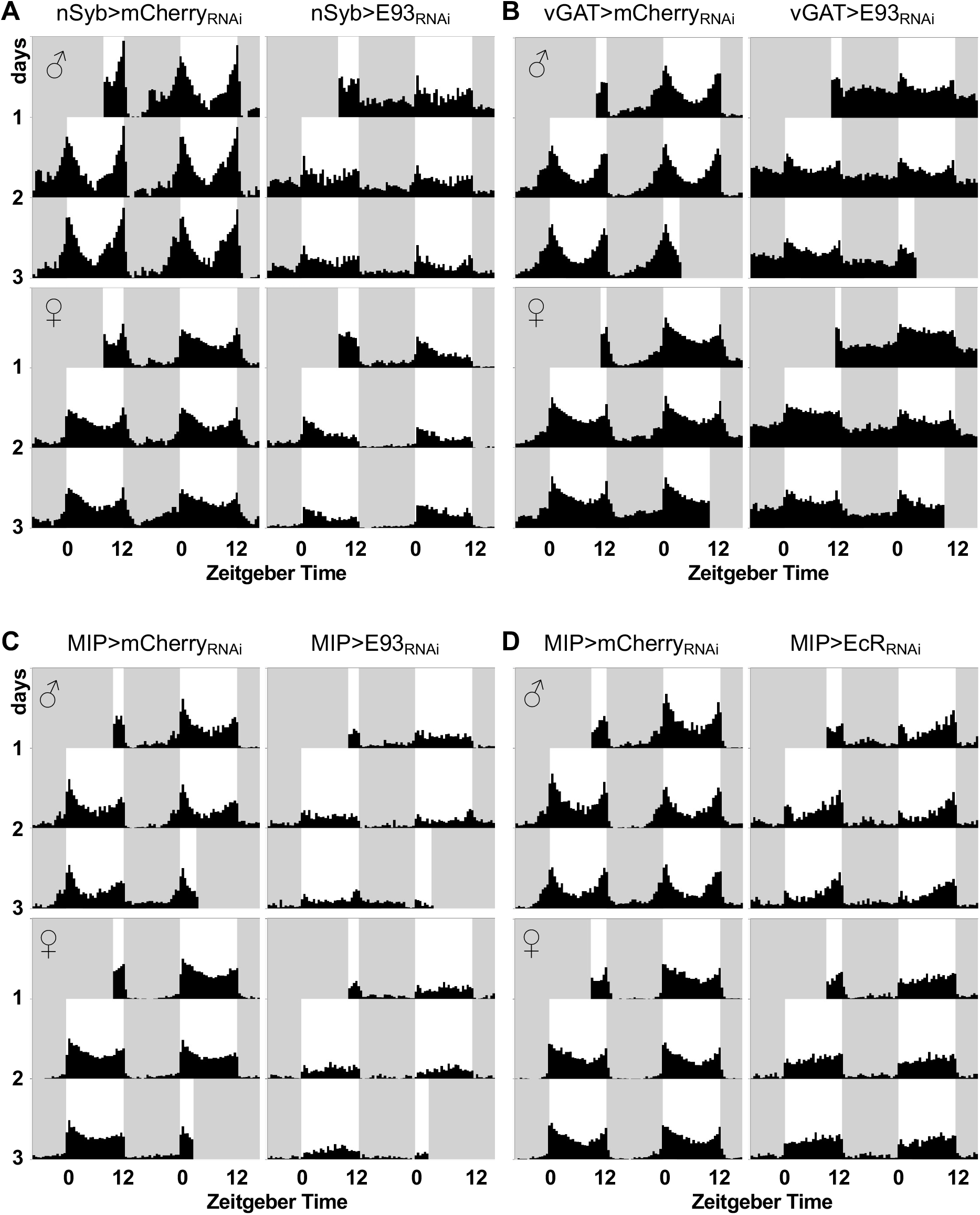
Knockdown of E93 attenuates morning and evening circadian anticipation. Beam breaks were recorded every 6 min from individual fly using the DAM assay system under 12 h light and 12 h dark conditions. Group average double-plotted actograms were generated with 30 min bins and plotted scale 0-120 counts / 30 min. The *x*-axis indicates Zeitgeber time and *y*-axis indicates days. Control (mCherry RNAi; left) and E93 or EcR knockdown (right) flies are plotted by sex (males (♂): top, females (♀): bottom). White background indicates light phase and gray background indicates dark phase. **A:** Pan-neuronal knockdown of E93 expression using the nSyb>GAL4 driver. nSyb>mCherry_RNAi_ males (N=27, 2 cohorts), nSyb>mCherry_RNAi_ females (N=29, 2 cohorts), nSyb>E93_RNAi_ males (N=31, 2 cohorts), and nSyb>E93_RNAi_ females (N=31, 2 cohorts) **B:** Knock down of E93 expression in GABA-ergic neurons. vGAT>mCherry_RNAi_ males (N=39, 3 cohorts), vGAT>mCherry_RNAi_ females (N=73, 5 cohorts), vGAT>E93_RNAi_ males (N=45, 3 cohorts), and vGAT>E93_RNAi_ females (N=64, 5 cohorts) **C:** Knock down of E93 expression in MIP neurons. MIP> mCherry_RNAi_ males (N=31, 2 cohorts), MIP> mCherry_RNAi_ females (N=32 2 cohorts), MIP> E93_RNAi_ males (N=32, 2 cohorts), and MIP> E93_RNAi_ females (N=32, 2 cohorts) **D:** Knock down of EcR expression in MIP neurons. MIP> mCherry_RNAi_ males (N=32, 2 cohorts), MIP> mCherry_RNAi_ females (N=28, 2 cohorts), MIP>EcR_RNAi_ males (N=30, 2 cohorts), and MIP>EcR_RNAi_ females (N=32, 2 cohorts)

Therefore, the increased nighttime activity observed in vGAT>E93_RNAi_ suggests that E93 could play a role in neuronal GABA activity to control food intake and promote sleep. Under standard 12 hour light and dark cycles, *Drosophila melanogaster* expresses a bimodal activity pattern with increased anticipatory activity prior to lights-on (dawn) and lights-off (dusk). Each anticipation is controlled by different subsets of circadian pacemaker neurons^57-59^. While control flies exhibit stereotypical anticipatory activities, these behaviors are almost absent in nSyb>E93 _RNAi_ flies (Figure 5A), suggesting that autonomous circadian clocks in knockdown flies are disrupted. In fact, in constant darkness, the majority of nSyb>E93 _RNAi_ flies show nearly arrhythmic activity patterns (Figure 6A), and only a few flies exhibit low amplitude of rhythms (1 out of 16 knockdown males, 2 out of 15 knockdown females, Figure S4A). Similar to nSyb>E93 _RNAi_ flies, vGAT>E93_RNAi_, MIP>E93_RNAi_, MIP>EcR_RNAi_ flies show moderate anticipation for lights-on or off (Figure 5B-D), suggesting that their circadian rhythms are also disrupted. vGAT>E93_RNAi_, MIP>E93_RNAi_, MIP>EcR_RNAi_ flies exhibited dampened circadian rhythms under constant darkness compared to that of controls. While nSyb>E93 _RNAi_, vGAT>E93_RNAi_, MIP>E93_RNAi_, MIP>EcR_RNAi_ flies all show disrupted circadian rhythm, the disruptions in nSyb>E93 _RNAi_ flies under constant darkness are the most severe (Figure 6B-D).

**Figure 6:**
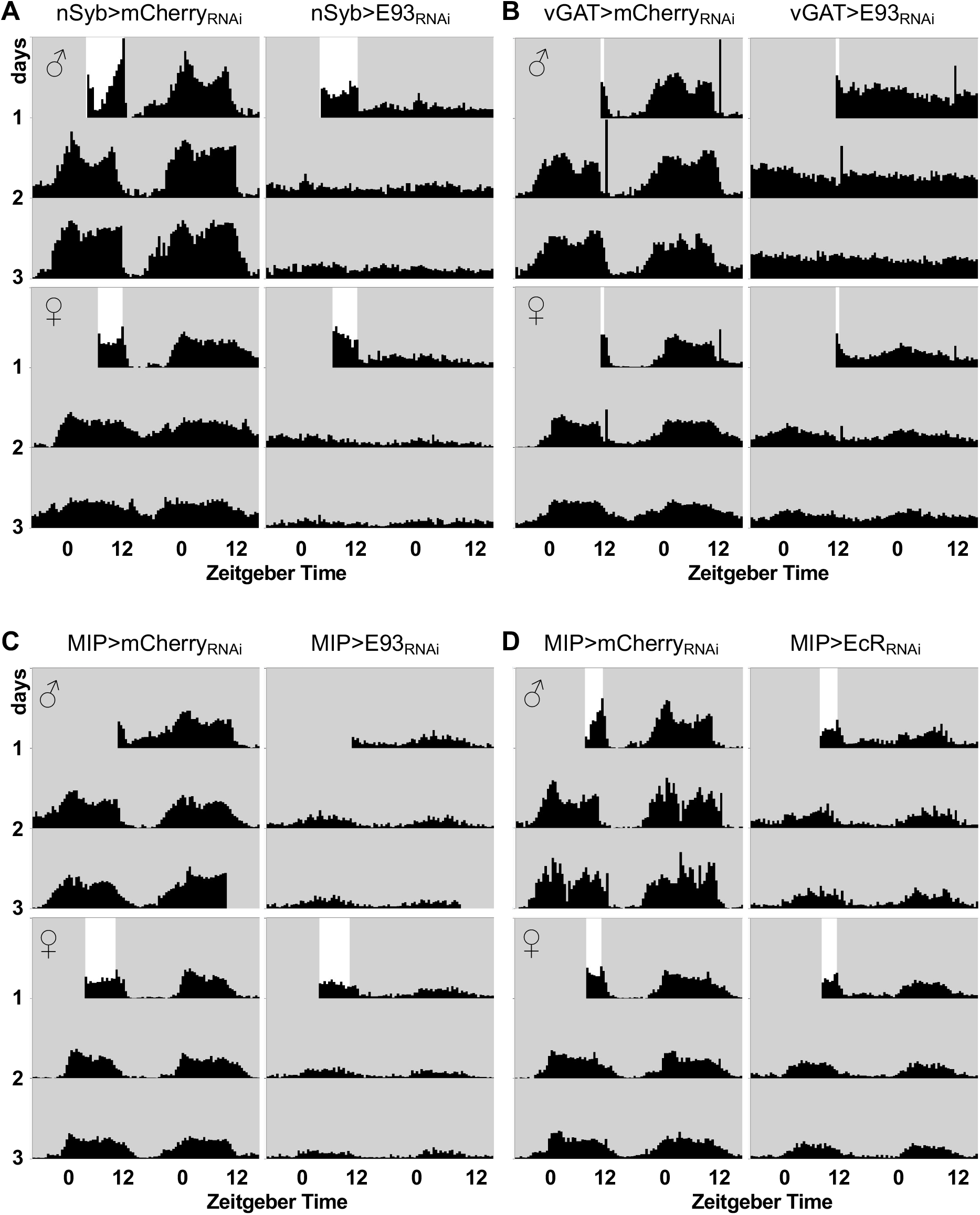
Reduction of E93 expression in neurons disrupts circadian rhythm. Group average double-plotted actograms of flies in constant darkness are shown. Flies raised in 12 hour light and 12 hour dark condition were placed in DAM during the light phase and released in constant darkness at the time of lights off. The white box indicates the last light cycle before releasing flies in constant darkness. **A:** Pan-neuronal knockdown of E93 expression using the nSyb>GAL4 driver. nSyb>mCherry_RNAi_ males (N=16, 1 cohort), nSyb>mCherry_RNAi_ females (N=15, 1 cohort), nSyb>E93_RNAi_ males (N=16, 1 cohort), and nSyb>E93_RNAi_ females (N=15, 1 cohort). **B:** Knock down of E93 expression in GABA-ergic neurons. vGAT>mCherry_RNAi_ males (N=26, 2 cohorts), vGAT>mCherry_RNAi_ females (N=39, 3 cohorts), vGAT>E93_RNAi_ males (N=31, 2 cohorts), and vGAT>E93_RNAi_ females (N=44, 3 cohorts). **C:** Knock down of E93 expression in MIP neurons. MIP> mCherry_RNAi_ males (N=31, 2 cohorts), MIP> mCherry_RNAi_ females (N=31 2 cohorts), MIP> E93_RNAi_ males (N=32, 2 cohorts), and MIP> E93_RNAi_ females (N=31, 2 cohorts). **D:** Knock down of EcR expression in MIP neurons. MIP> mCherry_RNAi_ males (N=31, 2 cohorts), MIP> mCherry_RNAi_ females (N=31, 2 cohorts), MIP>EcR_RNAi_ males (N=30, 2 cohorts), and MIP>EcR_RNAi_ females (N=32, 2 cohorts). Other conditions are the same as the actograms in Figure 5.

Together, our results suggest that E93 in the two subsets of neurons, MIP and vGAT, play an essential role in the regulation of metabolism, locomotive activity, and circadian rhythm.

## Discussion

In this study, we report a new role of E93, a conserved adult specifier^13^, in regulating feeding, metabolism, and biological rhythm. Neuron-specific knockdown of E93 causes multiple behavioral and physiological abnormalities such as incessant feeding, increased energy stores, and disrupted circadian rhythm. From screening of 17 different GAL4 drivers, we identified that E93 function in metabolism and circadian rhythm is primarily through its action in GABA-ergic and MIP neurons, both of which have been implicated in regulation of feeding, metabolism, and sleep^23,25,27,29,56^. Furthermore, we found that, E93 partly interact with the ecdysone receptor, EcR in MIP neurons. Although both males and females with neuron-specific knockdown of *E93* have increased energy stores, males favor storing excess energy as TAG whereas females preferentially store energy in the form of glycogen, suggesting a sexual dimorphism in energy storage. Since glycogen is the major resource accumulated in eggs, there might be an evolutionary advantage for females to store excess energy as glycogen rather than lipid. Increased frequency of feeding and resulting increased food intake most likely underlies the obesity in flies with neuron-specific knockdowns of *E93*.

Neuron-specific knockdown of *E93* also reduces activity and disrupts circadian rhythm. This is most pronounced when the flies are exposed to constant darkness (Figure 6). Similar circadian disruptions were observed in flies with GABA-specific or MIP-specific knockdowns of *E93*, supporting a role for E93 at the intersection of metabolism and circadian biology. All nSyb>E93_RNAi_, vGAT>E93_RNAi_, MIP>E93_RNAi_ and MIP>EcR_RNAi_ flies failed to show lights-on and off anticipatory behavior but exhibited either arrhythmicity or low amplitude rhythms under constant darkness. It is possible that reduced activity itself underlies the circadian phenotype. For instance, in a DAM assay, flies that stay near the food and spend a significantly longer time feeding than the controls would show decreased numbers of beam crosses. In our studies, vGAT>E93_RNAi_ flies had disrupted circadian rhythms, but activity levels were comparable to controls. Therefore, the constant darkness-induced arrhythmicity in males and low amplitudes in females is independent of activity. Taken together, the findings from vGAT>E93_RNAi_ and nSyb>E93_RNAi_ flies suggest that as least some part of disrupted circadian rhythm of nSyb>E93_RNAi_ is more likely associated with their metabolic abnormalities.

The mammalian homolog of E93 is LCOR (Ligand-dependent Corepressor) and has been shown to interact with nuclear receptors critical for metabolism^60-63^. This suggests that E93 might regulate circadian rhythm directly by interacting with the key nuclear receptors required for the metabolic clock^64,65^. Indeed, the fly orthologs of the key nuclear receptors, E75 (Retinoic acid-related Orphan Receptors in mammals) and DHR51 (Drosophila Hormone Receptor 51, REV-ERB in mammals) have conserved roles in regulating circadian rhythm^66,67^.

Although, the detailed neuro-molecular mechanisms by which E93 regulates these pleiotropic metabolic phenotypes require further investigation, we suggest that E93 is the key switch in shaping the adult brain to confer the fittest behavior, metabolism, and physiology in adults. As a transcription factor, E93 could directly control gene expression in coordination with specific nuclear receptors and switch the neuron’s profile from larva to adult, as suggested in Figure 7. Neuron-specific knockdown of *E93* could disrupt the function of several neuron types including GABA and MIP neurons. Our identification of GABA and MIP neurons as the major sites of action of E93 supports their roles as regulators of adult metabolism and physiology.

**Figure 7:**
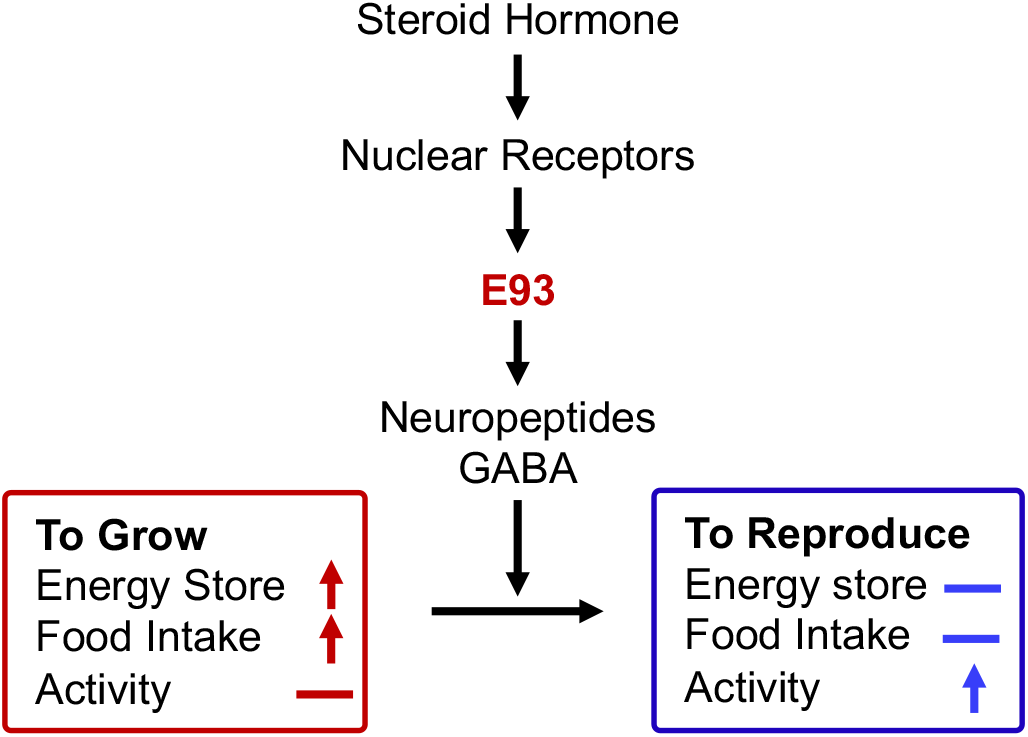
Model. E93 conveys a steroid signal to shape the nervous system optimized for adult metabolism and physiology.

A Recent study shows that several larvae-enriched genes are upregulated during metamorphosis in the E93 deletion mutant^18^. Interestingly, many of these genes are also implicated in neural fate determination (e.g. *sqeeze, ko*), metabolism (e.g. *secreted decoy of Inr*) and circadian rhythm (e.g. *clock*), supporting a role for E93 in the transition from larval to adult metabolism and physiology. One way by which E93 could control the transition from the larva brain to the adult brain is to establish the adult neuronal circuitry by controlling neurite pruning as its *C. elegans* ortholog does^14^. In fact, the nervous system undergoes dramatic rewiring often via steroid hormones during adolescence in many animals to switch the focus from growth to reproduction^68,69^. *Drosophila* also undergoes intensive remodeling of the nervous system during metamorphosis a stage in which E93 plays its essential roles^70^.

It is equally possible that E93 continues to function in GABA and MIP neurons in adults to maintain the function of adult neurons. In fact, several genes that play essential roles during development are repurposed to play roles in adults. For example, bursicon is essential for metamorphosis and also plays a role in energy homeostasis in adults^71^. That ecdysone, the upstream hormone of E93, continues to function in adults to control metabolism depending on reproductive state, suggests that E93 could continue to function downstream of the 20E steroid signal to control metabolic homeostasis in adults.

Together, our results provide insights into how a transcription factor acting in specific sets of neurons in the nervous system mediates steroid hormone signaling to shape and regulate adult metabolism and physiology.

## Author contribution

JE, SL and Y-JY planed and supervised the study. CY, SW, SY, Y-JY devised ideas, performed experiments, and analyzed the data. SY, CL and AR provided technical help, CY, SY and Y-JY wrote the manuscript with input from SW, AR, SL and JE.

## Acknowledgement

We thank Drs. Kim J, Fujikawa T, and Sieber M, and the Sieber lab members for invaluable discussions, Drs. Smith D and Kramer H for technical help, Drosophila Bloomington Stock Center, Korea Drosophila Resource Center, and Dr. Kim Y (GIST) for fly strains. This work was supported by the National Institutes of Health (NIH) through R01 DK100659 (JE), R01 NS114527 (SY), R01 AA019526 (AR), R01 AA026818 (AR), and by the National Science Foundation (NSF) through IOS-1931115 (SY).

There are no conflicts of interest.

