## Supplemental Materials for "Neuronal E93 regulates metabolic homeostasis"

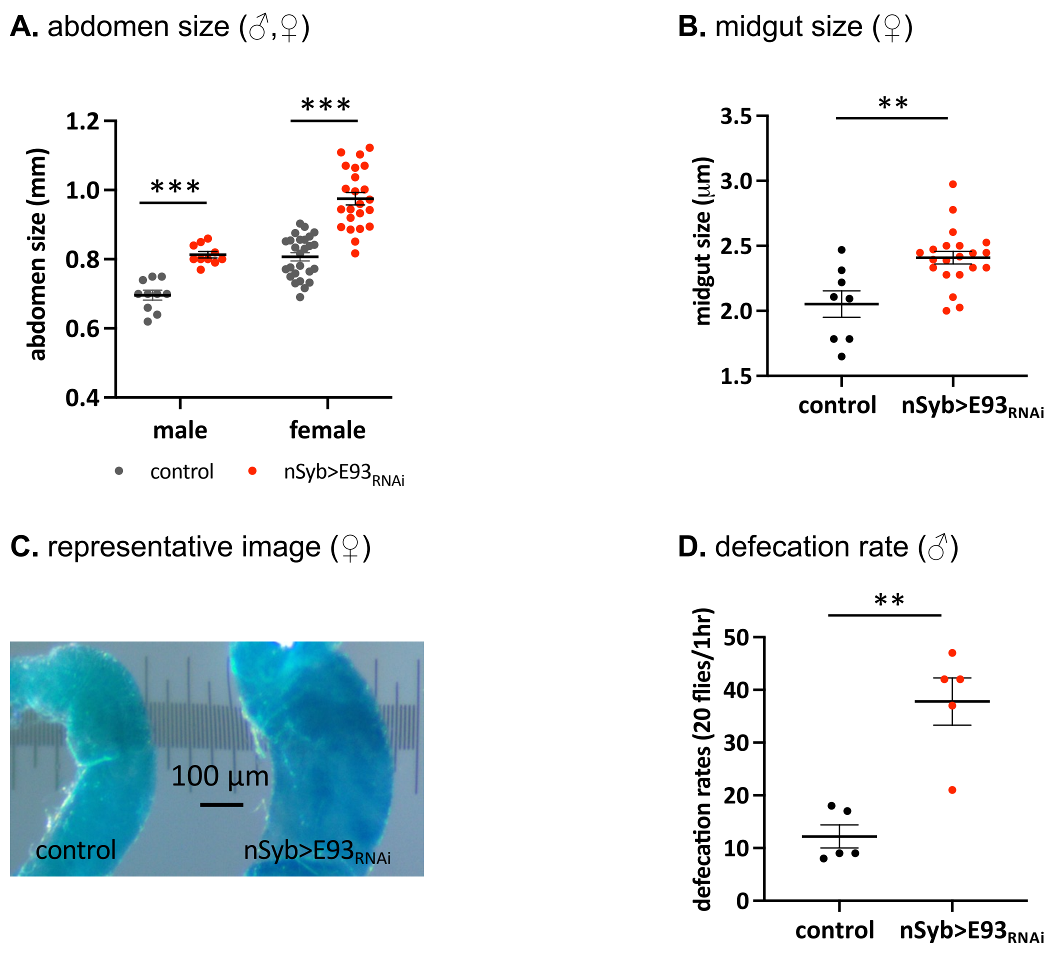


**Figure S1: Reduction of E93 expression in neurons increases abdomen size, midgut size and defecation rate.**

**A.** Reduction of neuronal E93 expression (nSyb>$E93_{RNAi}$) increases abdomen sizes in males and females (n=35 for nSyb>$E93_{RNAi}$ and n=32 for control).

**B.** Reduction of neuronal E93 expression (nSyb>$E93_{RNAi}$) increases the midgut size in females.

**C.** Representative image of midguts of a control and a nSyb>$E93_{RNAi}$ fly.

**D.** Defection rates increased in nSyb>$E93_{RNAi}$ males compared to controls. The data were plotted as the rate per 20 males per hour. The data was collected from five independent experiments, 20-40 males per genotype were used in each experiment.

A-B: Each data point represents one sample. The bars indicate the mean ± S.E.M. ** $p < 0.01$, *** $p < 0.001$.

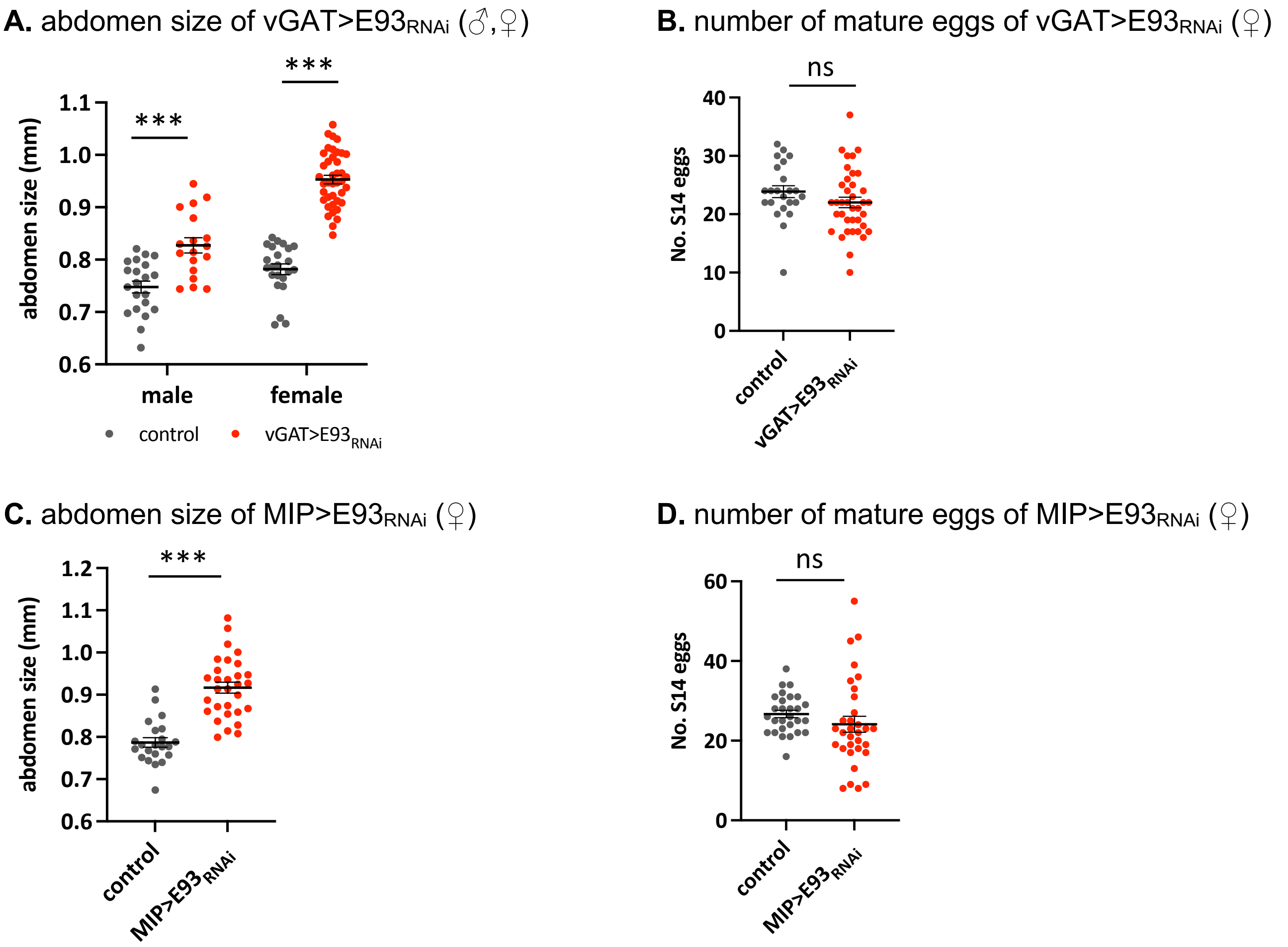


**Figure S2: Reduction of E93 expression specifically in GABA-ergic neurons or MIP-producing neurons increases abdomen size without an increase in the number of retained mature eggs.**

**A.** Reduction of E93 expression in GABA-ergic neurons (vGAT>$E93_{RNAi}$) increases abdomen sizes in males and females.

**B.** Reduction of E93 expression in GABA-ergic neurons does not change the number of retained mature eggs in females, suggesting that the increase in the abdomen size is not due to the increased number of the mature eggs.

**C-D.** Reduction of E93 expression in MIP-producing neurons (MIP>$E93_{RNAi}$) increases abdomen sizes in females (**C**) without changing the number of retained mature eggs in females (**D**).

Each data point represents one sample. The bars indicate the mean ± S.E.M.
*** $p < 0.001$, ns: not significant

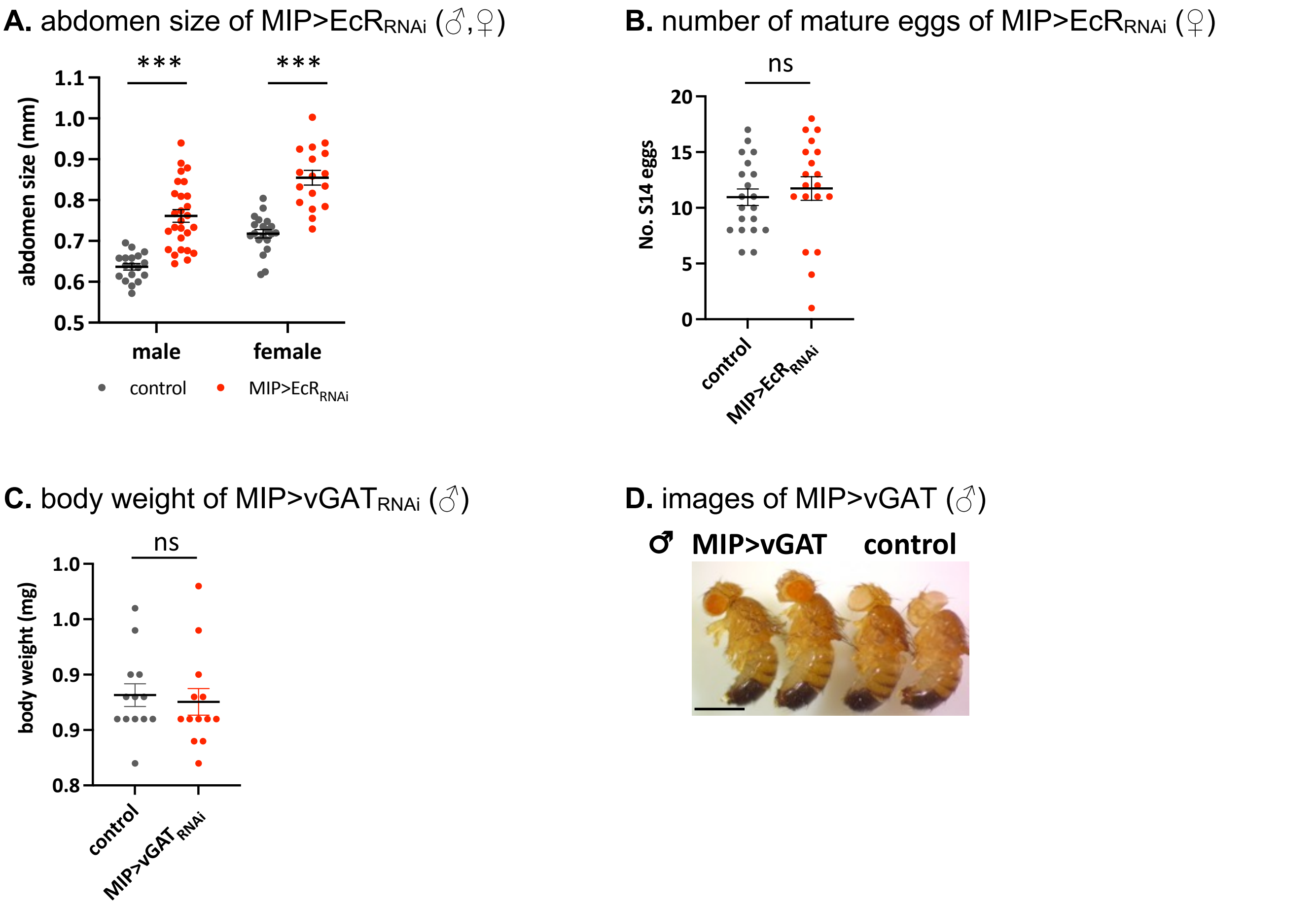


**Figure S3: Reduction of EcR (Ecdysone Receptor) expression specifically in MIP-producing neurons increases abdomen size without an increase in the number of retained mature eggs.**

**A.** Reduction of EcR expression in MIP neurons (MIP>$EcR_{RNAi}$) increases abdomen sizes in males and females.

**B.** Reduction of EcR expression in MIP neurons (MIP>$EcR_{RNAi}$) does not change the number of retained mature eggs.

**C.** Reduction of vGAT expression in MIP neurons (MIP>$vGAT_{RNAi}$) does not change the body weight.

**D.** Representative image of MIP>$vGAT_{RNAi}$ and the control (MIP>$mCherry_{RNAi}$). Scale bar = 1 mm

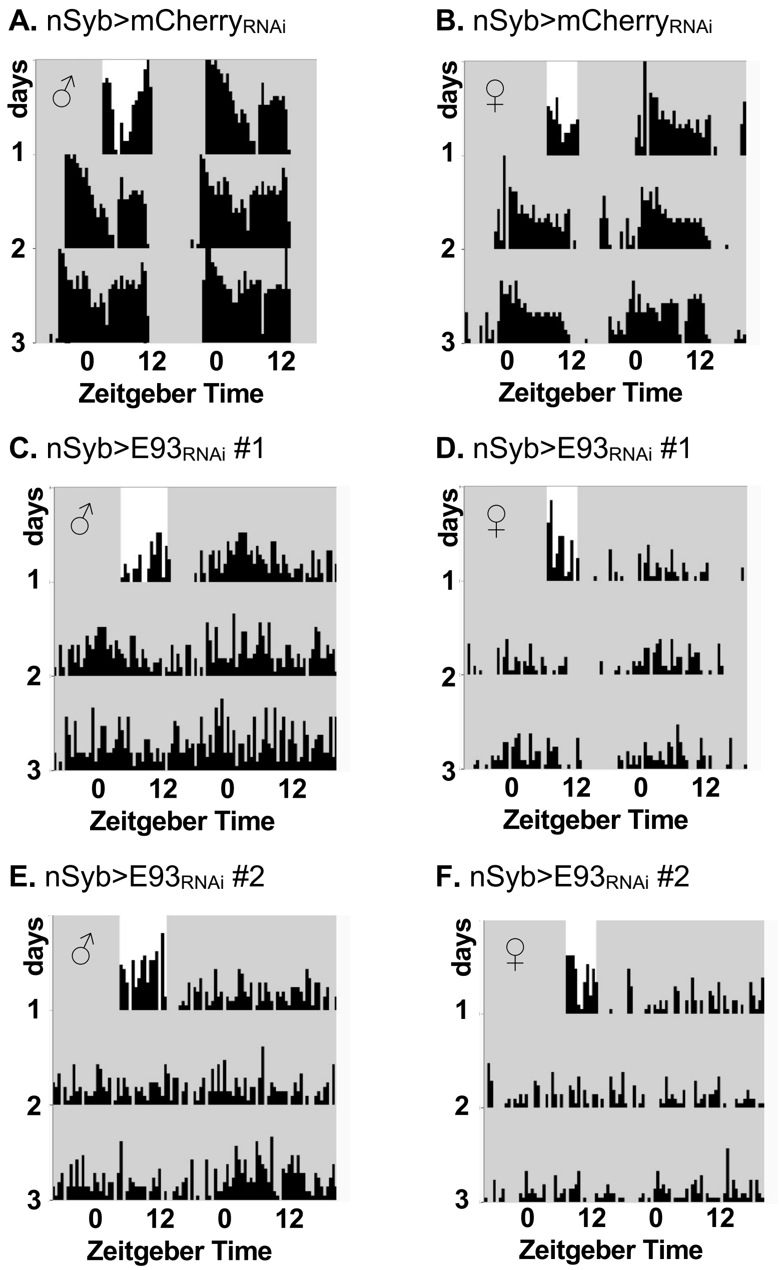


**Figure S4: Representative individual actograms of E93 knockdown flies in constant darkness.**

**A-B**: nSyb>mCherry$_{RNAi}$ male and female with intact rhythmicity.

**C-D**: nSyb>E93$_{RNAi}$ male and female with weak rhythmicity.

**E-F**: nSyb>E93$_{RNAi}$ male and female with arrhythmicity.

### Supplementary Methods

**Quantification of abdomen size.** Three-day old male and one and three-day old female flies of each genotype were incapacitated with dry ice, and then the wings and legs were removed to visualize the side of the abdomens. Flies were photographed and the width of the widest part was measured and calculated using a microscope calibration slide and Image J software.

**Quantification of midgut size.** The digestive tracts of female nSyb>$E93_{RNAi}$ and none>$E93_{RNAi}$ were dissected, and the midgut were identified by the position and the structure[1,2]. For the midgut, the width of the widest part was measured using a microscope stage calibration slide and Image J software. n=21 for nSyb>E93 females, n=8 for none>E93 females were used.

**Quantification of defecation.** Twenty male flies (three to five-days old) of each genotype were fed with food containing blue dye (0.1%) and transferred to 100 mm plates layered with filter paper[3]. After 60 min, the defecation spots were counted. The experiment was repeated five times.

**Quantification of matured eggs.** Five to ten virgin female flies were collected into a starvation vial (0.7% agar only) and fasted for two days. The flies were then placed on dry ice, their ovaries were dissected in 4% Paraformaldehyde (PFA, Fisher Scientific AA433689M), and the mature oocytes in stage 13-14 were counted based on opaqueness and size.

The *p* values were calculated by Mann-Whitney test for all quantifications of abdomen size, midgut size, defecation, and egg numbers.

### Supplementary videos

1. nSyb>E93$_{RNAi}$ flies remain on the food pad longer than the control.
https://1drv.ms/v/s!AjMFoQjl5fJOhhbSmunMXvHufISz?e=XzbVxb

2. Feeding video of the control flies (none>E93).
https://1drv.ms/v/s!AjMFoQjl5fJOhhmCAvUMMh0w_3-n?e=I9VM1c

3. Feeding video of nSyb>E93 flies.
https://1drv.ms/v/s!AjMFoQjl5fJOhhpFPx0uIe36q605?e=ZdZVRC
